# Glycoengineering of extracellular vesicles enhances cellular uptake and cargo delivery to target cells

**DOI:** 10.64898/2026.03.20.713142

**Authors:** Weihua Tian, Jiasi Chen, Anne Louise Blomberg, Judit Pina Agullet, Albert Juan Fuglsang-Madsen, Asha M. Rudjord-Levann, Helle Krogh Johansen, Søren Molin, Lasse Ebdrup Pedersen, Steffen Goletz

## Abstract

The glycocalyx is a major regulator of membrane recognition, yet its specific influence on extracellular vesicles (EVs) cellular uptake remains poorly defined. We established a genetic glycoengineering platform to systematically investigate how the major glycan classes on small EVs (sEVs) modulate cell interactions and functional cargo delivery. Using an isogenic panel of HEK293F lines lacking distinct glycan biosynthetic pathways, we find that removing glycosaminoglycans (ΔGAG-sEVs) yields a strong increase in cellular uptake and delivery of diverse cargos, including DNA oligonucleotides, siRNA, proteins, and plasmid DNA. Glycan-modified recipient cells show that sEV-cell communication and internalization is jointly governed by glycan features on both membranes. ΔGAG-sEVs strongly improve gene delivery and expression in recipient cells and in a physiologically relevant human airway epithelial model. These findings establish glycan structures as tunable regulators of sEV uptake and position ΔGAG-sEVs as potent vehicles for improved drug delivery and gene therapy.

## INTRODUCTION

Therapeutic delivery systems continue to face challenges related to bioavailability, biocompatibility, off-target effects, and limited efficiency in transporting and release of functional macromolecular cargos into target cells [1–3]. Extracellular vesicles (EVs), nano-sized particles released by most cell types, have emerged as promising delivery vehicles due to their native capacity to encapsulate and transfer biomolecules such as proteins, lipids, and nucleic acids [4–6]. Their roles in intercellular communication and diverse physiological and pathological processes have motivated increasing interest in harnessing EVs for drug and gene delivery applications [7–9]. However, improving EV cargo loading, ensuring efficient internalization into target cells and achieving effective intracellular delivery and endosomal release of functional macromolecules into the cyto- and nucleoplasm remain major obstacles to their therapeutic application [6,10–12].

Glycans are abundant on both cellular and EV surfaces and regulate essential biological processes, including cell adhesion, pathogen recognition, immune modulation, and biotherapeutic clearance [13–20]. The dense and compositionally diverse glycocalyx complicates systematic investigation, as glycan structures vary markedly across cell types, physiological states, and EV subpopulations [17,18,21–23]. EV heterogeneity further amplifies this complexity [23–25]. Although glycoengineering represents a promising strategy to dissect glycan function, existing approaches based on enzymatic deglycosylation have important limitations, such as incomplete and varying glycan removal, limited precision, and can affect protein stability or cause unintended biochemical changes [22,26–28]. Consequently, studies examining how specific glycan classes influence EV-cell interactions have yielded inconsistent conclusions, and no comprehensive analysis has addressed how major glycan classes on both EVs or recipient cells, alone or jointly, modulate vesicle-cell association and cargo transfer [22,23,29,30].

To overcome these limitations, we previously developed a cell-based glycoengineering platform, the “simple cell” strategy, to generate isogenic cell lines with defined glycan deficiencies by selectively disrupting individual glycan biosynthetic pathways [15]. sEVs produced from these cell lines exhibit correspondingly simplified glycan profiles, including the removal of complex N-glycans (ΔNG), O-GalNAc glycans (ΔOG), glycosphingolipids (ΔGSL), glycosaminoglycans (ΔGAG), or sialylation (ΔSA), thereby enabling controlled evaluation of individual glycan classes while minimizing biological heterogeneity [15].

In this study, we systematically investigate how these major glycan classes on both sEVs and recipient cells regulate vesicle-cell association and uptake, and the delivery of functional macromolecules. Using glycoengineered sEVs (GE-sEVs) loaded with different cargos either exogenously or through endogenous expression, we demonstrate glycan-dependent differences in cellular uptake and intracellular functional transfer of DNA oligonucleotides, siRNA, and plasmid DNA (graphic illustration shown in Fig. 1). Notably, GE-sEVs lacking glycosaminoglycans (ΔGAG-sEVs) displayed markedly enhanced cellular uptake and the highest efficiency in nucleic-acid transfer and transgene expression which includes effective endosomal escape, nuclear transport, transcription and translation. These findings identify glycan structures as tunable determinants of sEV-cell communication and establish ΔGAG-sEVs as promising platforms for improving and modulating the performance of sEV-based macromolecular delivery with a high potential for improved drug delivery and gene therapy.

**Fig. 1.**
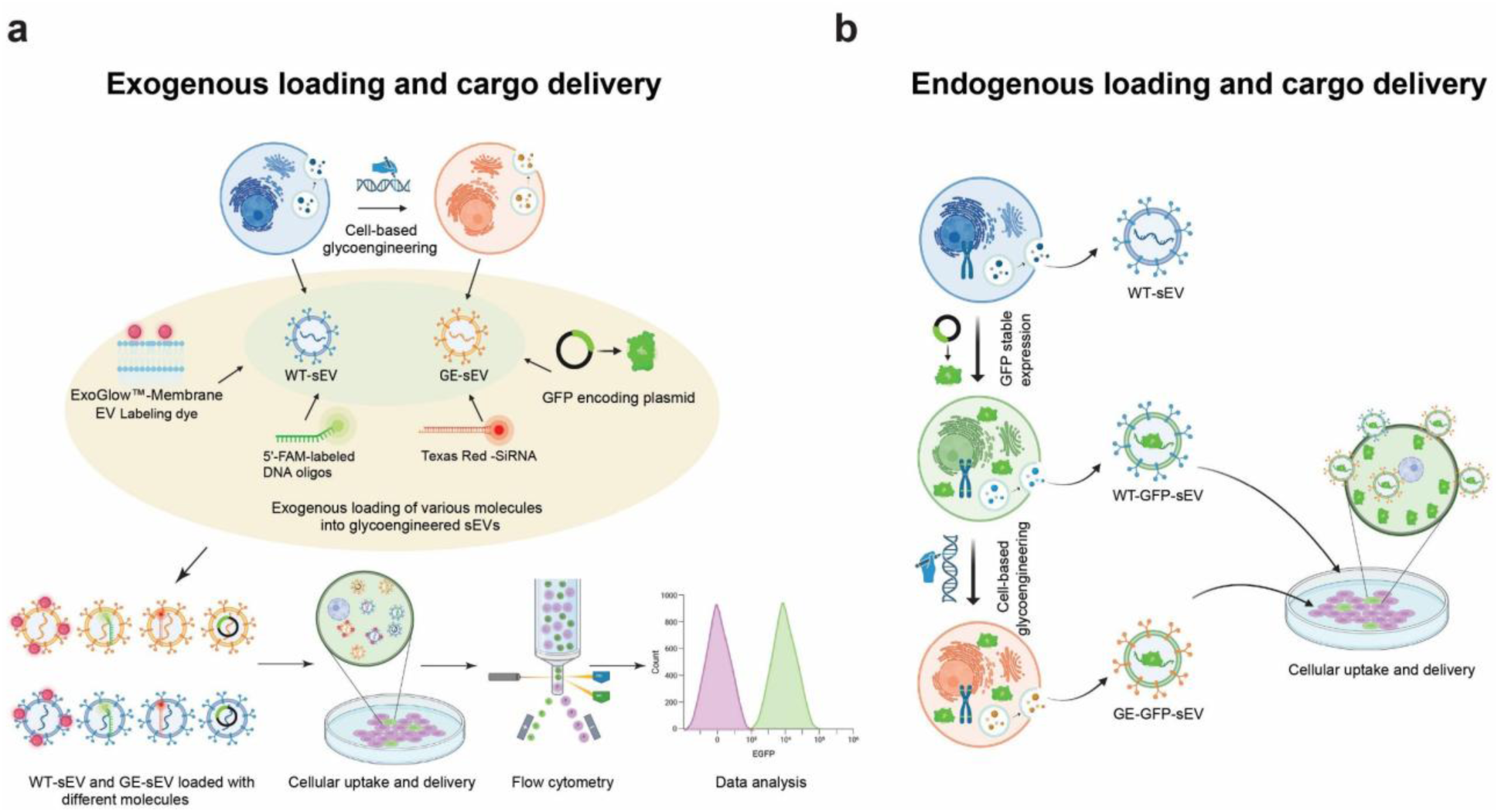
Overview of experimental strategies used to evaluate the impact of glycoengineering on sEV uptake and functional cargo delivery. **a,** Exogenous loading workflow: purified sEVs from WT and glycoengineered HEK293F cells were exogenously loaded with various molecules and applied to recipient cells for uptake assays. **b,** HEK293F WT cells were genetically engineered to stably express GFP, generating WT-GFP-sEVs; glycoengineering was then applied at the cellular level to produce GE-GFP-sEVs, and both vesicle types were used for uptake assays. Image partially created with BioRender.com.

## MATERIALS and METHODS

### Cell Culture

HEK293F cells were cultured as previously described [15]. Briefly, cells were cultured in FreeStyle 293 Expression Medium with 100 U/mL penicillin-streptomycin (Thermo Fisher Scientific = TFS) at 37 °C, 5% CO₂, in static culture flasks or 50 mL TPP TubeSpin Bioreactors (TPP Techno) under continuous shaking (150 rpm). Cells were passaged 2-3 times per week (seeding density 0.25×10^6^ cells/mL). NK92 cells were grown in Alpha MEM lacking ribonucleosides and deoxyribonucleosides with 0.2 mM inositol, 0.1 mM 2-mercaptoethanol, 0.02 mM folic acid, 100-200 U/mL recombinant IL-2, with 12.5% horse serum and 12.5% fetal bovine serum (FBS) and KHYG-1 cells in RPMI-1640 medium with 10% FBS and 100 U/mL penicillin-streptomycin (all TFS). Both NK92 and KHYG-1 cells were cultured at 37 °C, 5% CO₂ in static flasks.

### HEK293F sEV production and isolation

sEV production and purification were carried out as previously described [15]. Briefly, HEK293F cells were inoculated at 0.25×10^6^ cells/mL in FreeStyle 293 Expression Medium with 100 U/mL penicillin-streptomycin and 0.2% anti-clumping agent (TFS) and cultured at 37 °C, 5% CO₂ atmosphere with shaking at 150 rpm for 6 days. The medium was clarified by sequential centrifugation at 300×g for 10 min, 2,000×g for 10 min, and 10,000×g for 30 min to remove cells and debris. sEVs were then isolated by ultracentrifugation (considered the gold-standard method) twice at 100,000×g for 90 min, with a PBS wash in between to remove soluble proteins. The resulting sEV pellets were resuspended in PBS, aliquoted, and stored at −80 °C. All procedures were performed at 4 °C or on ice.

### sEV quantification through total protein determination

sEV quantification was performed as previously described [15]. Briefly, 2 μL of resuspended sEVs were mixed with 8 μL of RIPA lysis buffer (TFS), or 5 μL of resuspended sEVs was combined with 5 μL of 2×RIPA lysis buffer with EDTA-free cOmplete Protease Inhibitor Cocktail (Roche). The mixture was incubated on ice for 30 min to lyse the sEVs and release their protein content. Protein concentration was quantified using the Pierce BCA Protein Assay Kit (TFS), according to the manufacturer’s instructions, with BSA protein serving as the standard.

### sEV characterization using the Particle Metrix ZetaView

sEV particle size and concentration analyses were performed as previously described [15]. For zeta potential measurements, the ZetaView NTA system was used. sEV samples were diluted in 1% PBS (TFS) and measured in pulsed mode with three cycles, each scanning 11 positions. The captured videos were analyzed using ZetaView Software version 8.06.01.

### sEV labelling and uptake assay using ExoGlow

To label WT and glycoengineered sEVs, the ExoGlow-Membrane EV Labeling Kit (SBI) was used according to manufacturer’s instructions. 100 µg of purified sEVs in 400 µl PBS. To produce the labeling reaction buffer, 3 μl of labeling dye was blended with 18 μl of reaction buffer until the dye was entirely dissolved. 100 µg of purified sEVs in 400 µl PBS were mixed with the labeling dye and incubated at room temperature (RT) for 30 min. Unbound dye was removed by 4 rounds equilibration using PBS and centrifugation 800×g for 2 min using PD SpinTrap G-25 (Cytiva) and ExoQuick-TC precipitation. For each 100 μL of sample, 35 μL of ExoQuick-TC was added to the sEV, mixed, incubated on ice for 30 min, and centrifuged at 14,000×g for 10 min to pellet ExoGlow-labeled sEVs. As a negative control, PBS was incubated with labeling dye and treated identically to the sEV-containing samples. Finally, the ExoGlow-labeled sEVs pellet was resuspended in PBS and added to target cells. Cells were harvested at different time points by centrifugation, washed with 0.1% BSA/PBS, and resuspended in 100 µl IC Fixation buffer (TFS) for flow cytometry analysis using a MACS Quant 16 analyzer or visualized using the EVOS FL Auto 2 fluorescent microscope (TFS).

### sEV exogenous loading using Exo-Fect and cellular uptake assay

To load WT and glycoengineered sEVs with various nucleic acid molecules, the Exo-Fect Exosome Transfection Kit (SBI) was used according to the manufacturer’s instructions. For each 50 µg of sEVs, 10 µl of Exo-Fect solution were mixed with pMax-GFP plasmid DNA (Lonza), 5′-FAM-labeled DNA oligos (Macrogen), or Texas Red-conjugated siRNA (SBI) in PBS to a final volume of 150 µl. The transfection mixture was incubated at 37 °C shaking at 350 rpm for 10 min and then placed on ice. To terminate the reaction, 30 µl of ExoQuick-TC reagent was added, mixed by gentle inversion and pipetting, and incubated 30 min on ice. Samples were centrifuged 3 min at 14,000×g and the sEV pellet resuspended in culture medium and added to target cells. As a negative control, PBS was incubated with Exo-Fect solution and the corresponding molecules under identical conditions. Cells were harvested and analyzed as above using a MACSQuant 16 analyzer or EVOS FL Auto 2 fluorescence microscope.

### sEV endogenous loading of GFP, generation of GFP-sEV, and cellular uptake assay

HEK293F cells were transfected with the pMax-GFP plasmid using Lipofectamine 3000 (TFS) according to the manufacturer’s protocol. After four weeks of culture, GFP-positive cells were enriched and subjected to single-cell sorting using a MA900 cell sorter (Sony) to isolate individual clones. The clones were expanded for four weeks and one clone exhibiting stable and robust GFP expression selected for glycoengineering (designated HEK293F WT-GFP). Knockout of the *B4GALT7* gene was performed as previously described [15]. KO clones were validated for successful pheno- and genotypic knockout using fluorescent lectin staining to assess glycosylation changes and by Sanger sequencing. Three confirmed KO clones with comparable GFP expression were selected for further expansion and sEV production (designated HEK293F ΔGAG-GFP). sEVs were isolated and quantified as above [15]. For the cellular uptake assay, HEK293F WT or HEK293F ΔGAG cells (without GFP) were seeded at 10,000 cells per well in 100 µl of complete medium in 96-well plates. Purified GFP-sEVs derived from WT or ΔGAG clones were added at 15 µg or 35 µg per well. Cells were harvested and analyzed as above using a MACSQuant 16 analyzer.

### Air-Liquid Interface (ALI) lung epithelial model and sEV treatment

Basal cell immortalized-nonsmoker 1.1 (BCi-NS1.1) cells were cultured in Transwells at an air-liquid interface on 0.4-µm pore polyester membrane inserts (Corning), coated with type-I collagen (Gibco). Cells were seeded at 1.5×10^5^ cells/insert in expansion medium (PneumaCult-Ex Plus Medium, with PneumaCult-Ex Plus 50X (STEMCELL Technologies = ST) and further supplemented with 96 ng/mL hydrocortisone (ST) and 10 µM ROCK inhibitor (Y-27632,Tocris Bioscience), and maintained submerged until confluent (≈3 days), after which the apical medium was aspirated to establish ALI. Basolateral medium was aspirated, and ALI models were cultured in differentiation medium (PneumaCult-ALI Medium, with PneumaCult-ALI 10X Supplement and PneumaCult-ALI Maintenance Supplement (100X) (ST)) supplemented with 480 ng/mL hydrocortisone (ST) and 4 µg/mL heparin (ST) added to the basolateral compartment. Basolateral medium was refreshed twice weekly and ALI models were cultured for 28 days at 37 °C, 5% CO₂ until morphological and transepithelial electrical resistance (TEER) criteria for differentiation were met. sEV preparations were exogenously loaded with pMax-GFP plasmid Exo-Fect as described above and applied to the apical surface in 10 µl PBS to simulate inhalational exposure. Two doses of sEV were tested per experiment: 15 µg and 60 µg. Untreated (PBS alone) ALI models served as negative controls. After application, ALI models were incubated at 37 °C, 5% CO₂ for 24 h.

### Immunofluorescent staining of ALI models

ALI models were stained for F-actin using Phalloidin-AlexaFluor 647 (TFS). Fluorescence imaging was performed with excitation at 650 nm and emission detected at 665-690 nm for F-actin. Nuclei were counterstained with DAPI (TFS). ALI models were fixed in 4% paraformaldehyde for 20 min at 4 °C, washed 3 times in PBS, and incubated in blocking buffer (0.1% Triton X-100, 1% saponin, 3% BSA in PBS) for 1 h at RT. Next, ALI models were incubated with the dyes diluted in staining solution (3% BSA and 1% saponin in PBS) for 2 h at RT. After staining, membranes were excised from the Transwells and mounted onto microscope slides with VECTASHIELD Antifade Mounting Medium (VWR). Images were acquired with a Leica Stellaris 8 Confocal Microscope.

### Microscopy and image analysis

GFP expression was evaluated by confocal light microscopy on a Leica Stellaris 8 up-right microscope with identical exposure settings across conditions; excitation at 488 nm, emission detected at 501-540 nm, for GFP. GFP- and DAPI fluorescence integrated densities were quantified using the Fiji/ImageJ software tool “3D Object Counter”. ZStacks were analyzed as 16-Bit (Image > Type > 16-Bit) and were split into blue (epithelial nuclei stained with DAPI), red (epithelial F-actin, stained by phalloidin-Alexa), and green channels (GFP expressed by BCi cells). When brightness/contrast adjustments or noise subtraction were required, the same processing parameters were applied to both fluorescence channels (DAPI and GFP) before analyzing individual ZStacks. Some ZStacks did not require any preprocessing. In cases where noise reduction was necessary, pixel values were reduced using Fiji (Mac) via Process > Math > Subtract. For the 3D Object Counter analyses, measurements were configured to report Volume, Integrated Density, Centroid, and Mean Gray Value, with a threshold value set at 10,000. Objects above 1 µm^3^ were deemed as true signal for both DAPI and GFP channels. The reported Delivery-& Translation Coefficients (DTC) are reported as the total GFP integrated density (pixel intensity units×pixel area) divided by the total DAPI integrated density to normalize the GFP integrated density to the number of epithelial cells present in the given ZStack (see equation 1).

Equation 1. DTC, which is the total GFP integrated density per thousands of total DAPI integrated density.

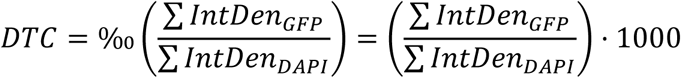

### Statistics

All statistical analyses were performed using GraphPad Prism software. Data are reported as mean ± standard deviation (s.d.). Group differences were assessed using two-way ANOVA with Dunnett’s or Sidak’s multiple-comparison tests, as specified in the figure legends. The number of replicates for each experiment is provided in the corresponding legends.

## RESULTS

### Modification of sEV glycosylation strongly modulates cellular uptake

To assess how specific glycan classes on sEVs influence interactions with recipient cells, we quantified uptake of WT and five glycoengineered sEV variants through exogenously loading an EV specific membrane dye. The sEV variants were produced from isogenic HEK293F lines with selective disruptions in major glycosylation pathways: ΔNG (*MGAT1* KO), ΔOG (*C1GALT1C1* KO), ΔGSL (*B4GALT5/6* KO), ΔSA (*GNE* KO), and ΔGAG (*B4GALT7* KO), as detailed in our previous publication [15]. Assay conditions were optimized, and two doses (0.78 μg and 1.56 μg) and two time points (7 h and 20 h) were selected for analysis. To minimize potential effects of clonal variability, sEVs from three independent clones of each variant and three separate batches of HEK293F WT cells were pooled for uptake quantification. Glycan modification produced strikingly divergent uptake phenotypes. ΔSA-sEVs displayed significantly enhanced uptake at both 7 h (Fig. 2a) and 20 h (Fig. 2b), as reflected by a marked increase in Mean Fluorescence Intensity (MFI), indicating improved delivery efficiency upon desialylation. Among all variants, ΔGAG-sEVs exhibited the strongest enhancement relative to WT across all doses and time points (Fig. 2a and 2b). In contrast, other glycan modifications produced only modest or non-consistent effects on uptake, underscoring the dominant inhibitory role of GAGs in sEV-cell association.

**Fig. 2.**
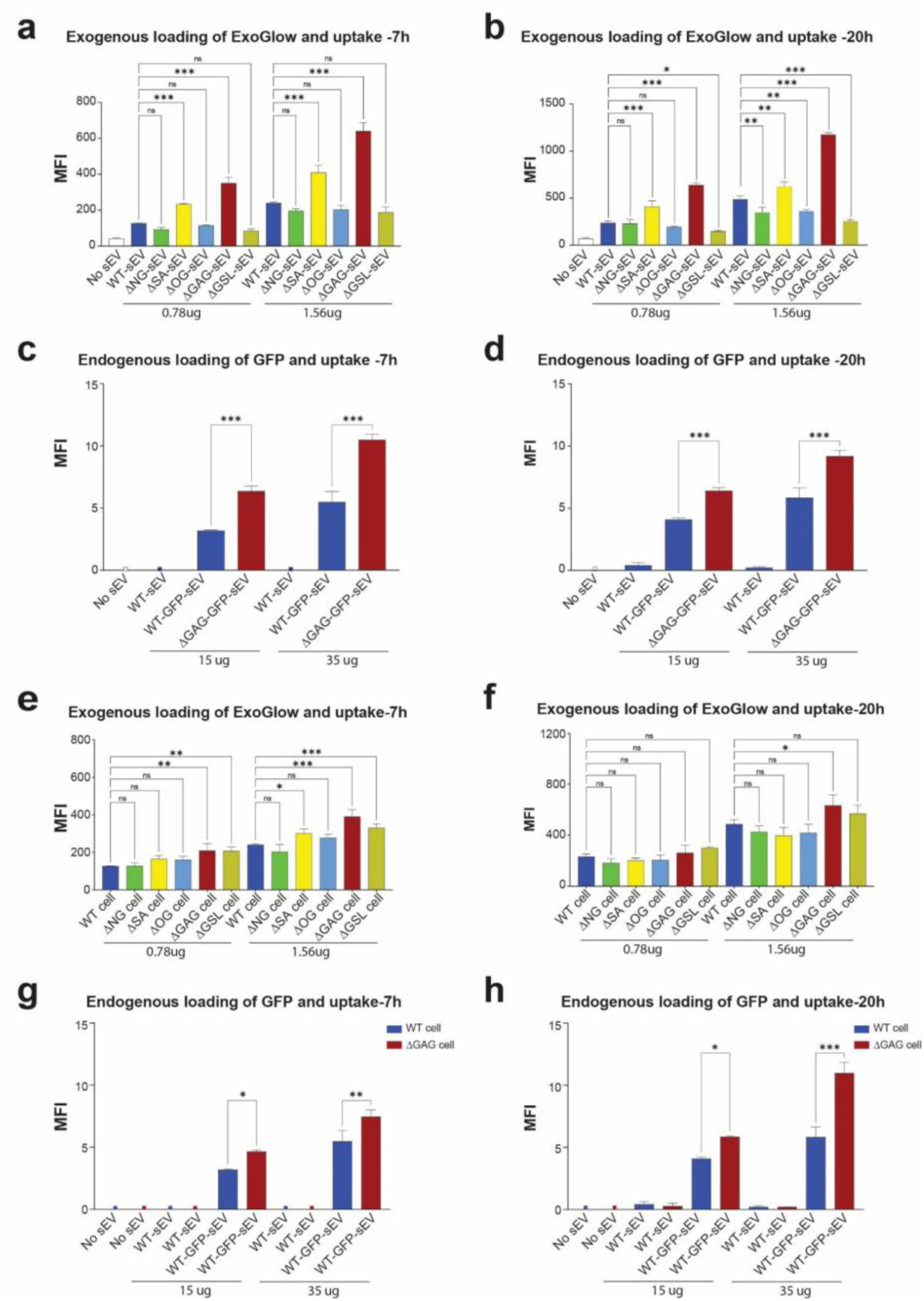
Vesicle and cellular glycans govern sEV uptake in HEK293F cells. **a, b,** Uptake of six sEV variants—WT, ΔNG, ΔSA, ΔOG, ΔGAG, and ΔGSL—by HEK293F WT cells was quantified using ExoGlow-Membrane-labeled sEVs. Cells were incubated with 0.78 µg or 1.56 µg sEVs, and mean fluorescence intensity (MFI) was measured by flow cytometry after 7 h (**a**) and 20 h (**b**). **c,d,** Validation using endogenously GFP-loaded sEVs. HEK293F WT recipient cells were incubated with 15 µg or 35 µg WT-GFP-sEVs or ΔGAG-GFP-sEVs. MFI was assessed at 7 h (**c**) and 20 h (**d**). **e,f,** Uptake of WT-sEVs was examined in HEK293F WT cells and five isogenic glycoengineered knockout lines lacking complex N-glycans (ΔNG), sialic acids (ΔSA), extended mucin-type O-glycans (ΔOG), glycosaminoglycans (ΔGAG), or glycosphingolipids (ΔGSL). ExoGlow-Membrane-labeled sEVs (0.78 µg or 1.56 µg) were applied, and MFI was measured at 7 h (**e**) and 20 h (**f**). **g,h,** Uptake of endogenously loaded WT-GFP-sEVs was compared between WT and ΔGAG recipient cells using 15 µg or 35 µg sEVs. MFI was measured at 7 h (**g**) and 20 h (**h**). Data are presented as mean ± s.d. from three replicates. P values were calculated using two-way ANOVA with Dunnett’s multiple-comparison tests for panels **a, b, e, and f**, and Sidak’s multiple-comparison tests for panels **c, d, g, and h**: *P < 0.05; **P < 0.01; *P < 0.001.

### Endogenously GFP-loaded sEVs validate the ΔGAG uptake phenotype

To further validate the role of GAGs in sEV uptake, we generated HEK293F cell lines stably expressing GFP, enabling endogenous loading of the GFP fluorescent protein into secreted sEVs (hereafter WT-GFP-sEV). We then introduced *B4GALT7* knockout into this the GFP-stably expressing cell line to generate glycoengineered sEVs lacking glycosaminoglycans (hereafter ΔGAG-GFP-sEV). Resulting WT-GFP-sEVs and ΔGAG-GFP-sEVs contained comparable amounts of GFP. Recipient HEK293F WT cells were incubated with either 15 µg or 35 µg of GFP-sEVs, and flow cytometry MFI was quantified at 7 h and 20 h (Fig. 2c and 2d). Across both doses and time points, ΔGAG-GFP-sEVs showed markedly higher cellular uptake than WT-GFP-sEVs, fully mirroring the trends observed with dye-labeled sEVs. These data demonstrate that enhanced uptake of ΔGAG-sEVs is an intrinsic property of GAG removal and is independent of external loading and the cargo-labeling strategy.

### Recipient-cell glycans also modulate sEV uptake

We next examined whether glycans on recipient cells influence sEV uptake. WT-sEVs derived from HEK293F cells were exogenously loaded with the dye and applied to a panel of glycoengineered HEK293F lines. To minimize clonal variability, three independent clones were analyzed for each glycan-modified variant. Uptake of WT-sEVs into these cells revealed that loss of complex N-glycans (ΔNG) or extended mucin-type O-glycans (ΔOG) on cells had minimal impact. In contrast, cells lacking GSLs (ΔGSL) or GAGs (ΔGAG) exhibited markedly increased sEV uptake (Fig. 2e and 2f), indicating that these glycans act as steric or electrostatic barriers to vesicle engagement. To validate these findings with endogenously loaded sEV, we performed parallel experiments using WT-GFP-sEVs. WT-GFP-sEVs were applied to WT and ΔGAG recipient cells at 15 µg and 35 µg dose, and cellular uptake was quantified at 7 h and 20 h (Fig. 2g and 2h). ΔGAG cells consistently displayed higher uptake, confirming that cell-surface GAG function as bidirectional regulators that suppress sEV entry.

### Matrix analysis reveals coordinated glycan-glycan determinants of sEV uptake

To dissect combinatorial glycan effects, we performed a matrix analysis testing all sEV glycovariants across all glycoengineered recipient cells. Overall, most glycoengineered cell types exhibited uptake levels comparable to WT; however, ΔGAG-sEVs consistently showed the highest uptake in all cells across conditions, followed by ΔSA-sEVs (Fig. 3a-3d). Among recipient-cell variants, ΔGSL cells, and often also ΔGAG cells, displayed notably increased uptake across multiple GE-sEV types, indicating that both GSLs and GAGs on the cell surface can restrict sEV uptake. At the 7 h but not 20h time point, ΔSA cells also showed improved uptake, suggesting dose- and time-dependent modulation by sialylation. Importantly, ΔGAG-sEVs did not achieve maximal uptake in ΔGAG recipient cells. Instead, ΔGSL and ΔSA cells supported the strongest uptake for ΔGAG-sEVs, demonstrating that optimal vesicle-cell interactions arise from specific glycan complements rather than matching deficiencies. Parallel experiments using GFP-sEVs confirmed these trends and further showed that removal of GAGs on either vesicles or cells enhances uptake (Fig. 3e and 3f). Collectively, these findings reveal that glycan-dependent vesicle engagement follows coordinated, non-linear rules that can only be resolved using a clean isogenic glycoengineering framework. The results highlight the complexity of glycan-glycan interactions and underscore the need for systematic glycoengineering to rationally optimize sEV-cell communication.

**Fig. 3.**
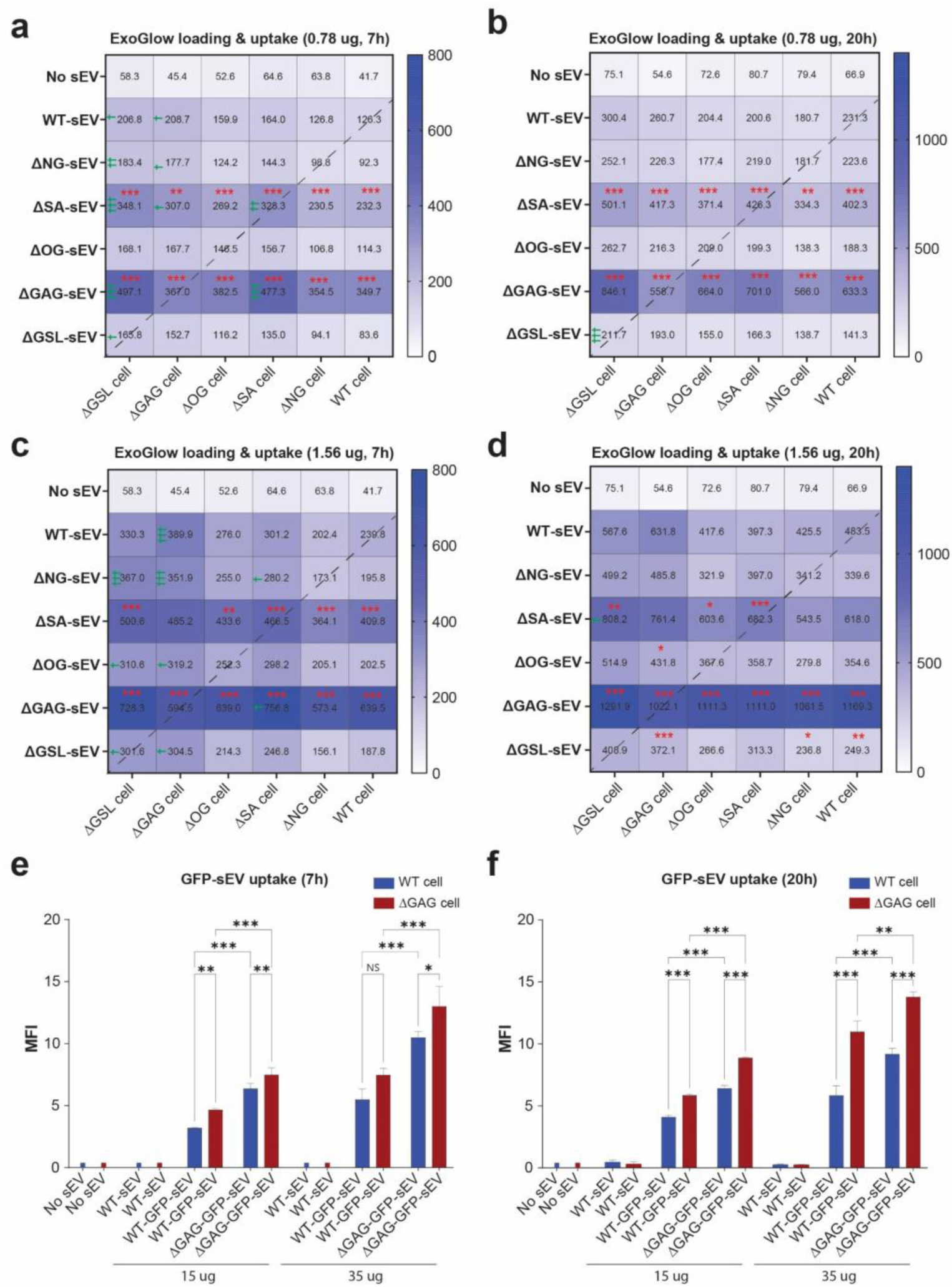
Matrix analysis of EV-cell glycan interactions. **a-d,** Pairwise uptake analysis between six sEV variants (WT, ΔNG, ΔSA, ΔOG, ΔGAG, ΔGSL) and six isogenic HEK293F recipient cell lines (WT, ΔNG, ΔSA, ΔOG, ΔGAG, ΔGSL). Heatmaps display mean fluorescence intensity (MFI) under four conditions: (**a**) 7 h, 0.78 µg sEV; (**b**) 20 h, 0.78 µg sEV; (**c**) 7 h, 1.56 µg sEV; (**d**) 20 h, 1.56 µg sEV. Values represent the mean MFI from three independent clones per glycoengineered condition. Statistical comparisons: Asterisks (* red): sEV variants compared to WT-sEV within the same recipient cell type. Daggers († green): uptake of the same sEV variant compared between glycoengineered and WT recipient cells. **e,f,** Validation using endogenously GFP-loaded sEVs. WT and ΔGAG recipient cells were treated with 15 µg or 35 µg WT-GFP-sEVs or ΔGAG-GFP-sEVs. MFI was quantified at 7 h (**e**) and 20 h (**f**). Data are presented as mean ± s.d. from three replicates. P values were calculated using two-way ANOVA with Sidak’s multiple-comparison tests: */† P < 0.05; **/†† P < 0.01; ***/††† P < 0.001.

### HEK293F ΔGAG-sEVs enhance uptake in clinically relevant NK cell lines

To evaluate whether glycan-dependent uptake extends beyond HEK293F cells, we assessed sEV internalization in NK-92 and KHYG-1 cells—two clinically relevant, hard-to-transfect suspension cell lines. Uptake trends observed in HEK293F cells were maintained: ΔGAG-sEVs consistently showed the highest uptake with a 5.5-to-7.5 fold increase for all 3 recipient cell lines in comparison to WT-sEV after 7h and 20 h, respectively, followed by ΔSA-sEVs with an according 2 to 2.5 fold increase for all 3 cell lines (Fig. 4a and 4b). Despite identical sEV dosing, the overall uptake into NK92 was a bit lower than that into KHYG-1 and HEK293F cells. This confirms that the ΔGAG enhancement effect is robust and applies for diverse target cells.

**Fig. 4.**
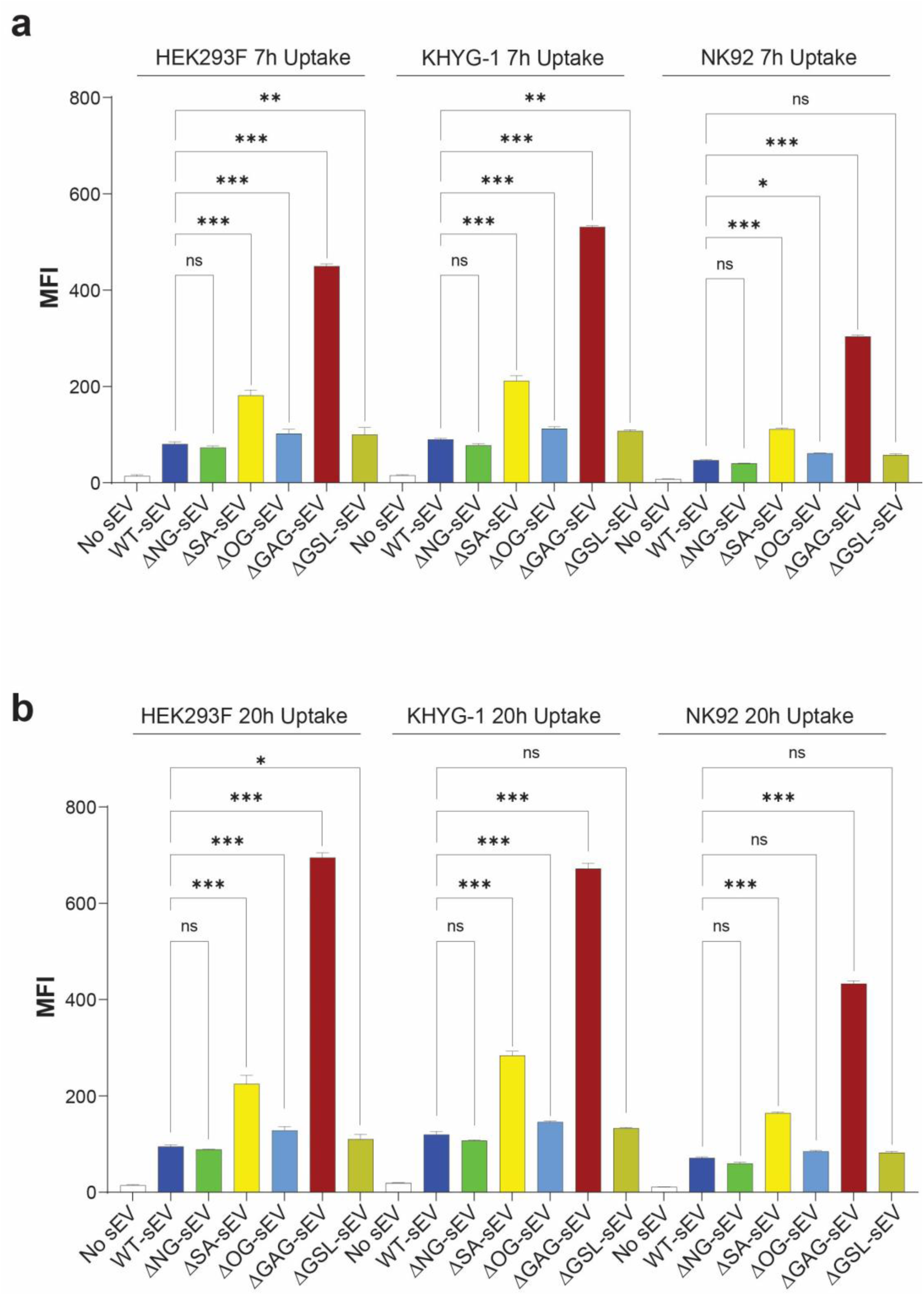
Uptake of HEK293F GE-sEVs by HEK293F, KHYG-1, and NK92 Cells. Uptake of six HEK293F-derived sEV variants (WT, ΔNG, ΔSA, ΔOG, ΔGAG, ΔGSL) by HEK293F, KHYG-1, and NK92 cell lines was assessed using ExoGlow-Membrane-labeled sEVs. **a,** MFI measured at 7 h after incubation with 1.56 µg sEVs. **b,** MFI measured at 20 h using the same dose. Data are presented as mean ± s.d. from three replicates. P values were calculated using two-way ANOVA with Dunnett’s multiple-comparison tests: *P < 0.05; **P < 0.01; ***P < 0.001.

### Glycoengineered sEVs enhance delivery of functional nucleic acids

To determine whether improved uptake results in enhanced functional transfer of therapeutic macromolecules, we evaluated the delivery of FAM-labeled DNA oligonucleotides and Texas Red-conjugated siRNA. These fluorescently labeled cargos represent distinct therapeutic modalities and allow quantitative assessment of intracellular accumulation by flow cytometry. ΔGAG-sEVs delivered FAM-labeled DNA oligonucleotides with the highest efficiency, increasing the fraction of FAM⁺ cells from 5.24% with WT-sEVs to 14.2% with ΔGAG-sEVs (Fig. 5a). Delivery of Texas Red-siRNA followed a similar pattern. While overall gating yielded ∼20% Texas Red⁺ cells across conditions, applying a stringent high-intensity gate (TXred⁺⁺) revealed a clear improvement from 0.30% with WT to 2.03% with ΔGAG-sEVs. ΔSA sEVs provided intermediate enhancement, whereas ΔNG-sEVs generally reduced transfer (Fig. 5b). Together, these results demonstrate that sEV glycan modification, particularly removal of GAGs, can substantially enhance their capacity as carriers for nucleic acid therapeutics.

**Fig. 5.**
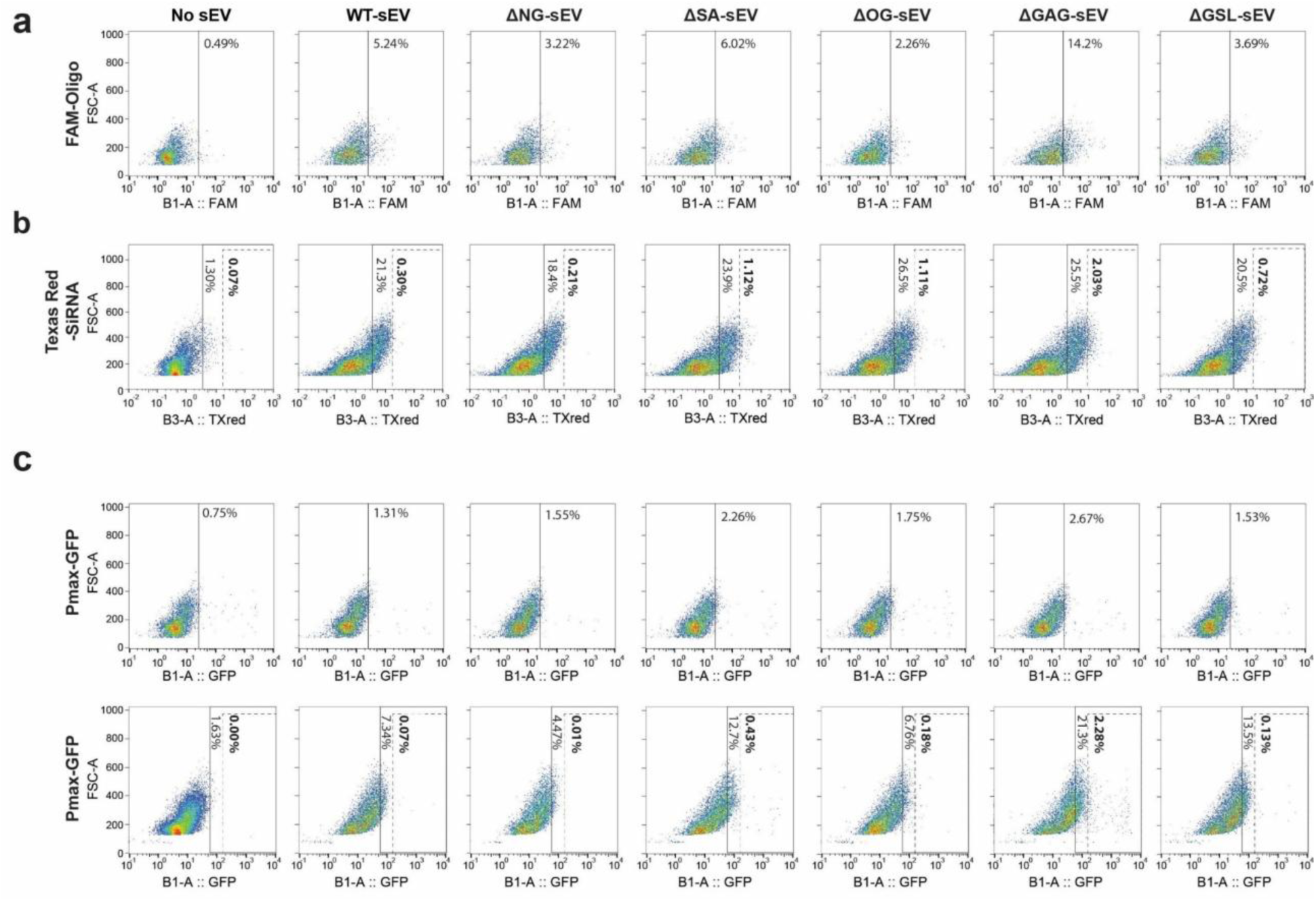
Functional delivery of therapeutic macromolecules using glycoengineered sEVs. Functional delivery of three classes of cargo via WT and glycoengineered sEVs into HEK293F WT cells was assessed by flow cytometry 20 h post-delivery: **a,** 12.5 µg sEVs loaded with 5 pmol FAM-labeled DNA oligonucleotides. **b,** 25 µg sEVs loaded with 1.25 pmol Texas Red-conjugated siRNA. **c,** 12.5 µg sEVs loaded with 1.25 µg pMax-GFP plasmid DNA (upper panel) or 25 µg sEVs loaded with 2.5 µg pMax-GFP plasmid DNA (lower panel). Uptake or functional expression was quantified by flow cytometry, and representative dot plots are shown.

### Glycoengineered sEVs improve plasmid DNA delivery and transgene expression

To evaluate delivery of larger macromolecules, we analyzed transfer of a GFP-encoding plasmid. Plasmid delivery requires cellular uptake, endosomal escape, nuclear localization, and subsequent GFP expression through transcription and translation, making it a representative model for gene-replacement or gene-addition therapies. GFP expression in HEK293F WT cells was assessed by flow cytometry using two doses of sEVs. At the low sEV dose, ΔGAG-sEVs produced a modest but consistent increase in GFP⁺ cells, rising from 1.31% for WT-sEVs to 2.67% (Fig. 5c upper panel). At the higher dose, this enhancement became substantial, with GFP⁺ cells increasing 3-fold from 7.34% to 21.3%. When applying a stringent strong GFP (GFP⁺⁺) gate to capture only strongly expressing cells, ΔGAG-sEVs yielded a >30-fold improvement, increasing from 0.07% to 2.28% (Fig. 5c lower panel). Importantly, the stringent GFP⁺⁺ gate likely enriches for cells that have undergone more efficient cellular uptake and nuclear delivery, highlighting a functional advantage. Together, these results indicate that glycan modification enhances not only vesicle binding, but also downstream intracellular processes required for efficient plasmid delivery and transgene expression.

### ΔGAG-sEVs enhance gene delivery in an airway air-liquid interface model

To evaluate translational potential, we used a differentiated human airway epithelium air-liquid interface model which mimics the lung barrier with secreted mucus and cilia movement and enables testing of apical sEV delivery. WT-sEV and ΔGAG-sEVs preloaded with pMax-GFP plasmid were applied to the apical surface to model inhalation-based administration and GFP expression was analyzed and quantified 24 h post-application by confocal microscopy. GFP expression was ∼15-fold higher in ΔGAG-sEV-treated cultures (Fig. 6a and 6b), demonstrating superior plasmid delivery and gene expression across a physiologically relevant epithelial barrier. All groups maintained stable transepithelial electrical resistance (TEER) values, indicating that neither WT nor ΔGAG-sEV treatment meaningfully affected epithelial integrity or compromised barrier function (Fig. 6c). Together, these results show that ΔGAG-sEVs substantially enhance plasmid DNA delivery and transgene expression in a representative human airway epithelial model while preserving epithelial barrier integrity.

**Fig. 6:**
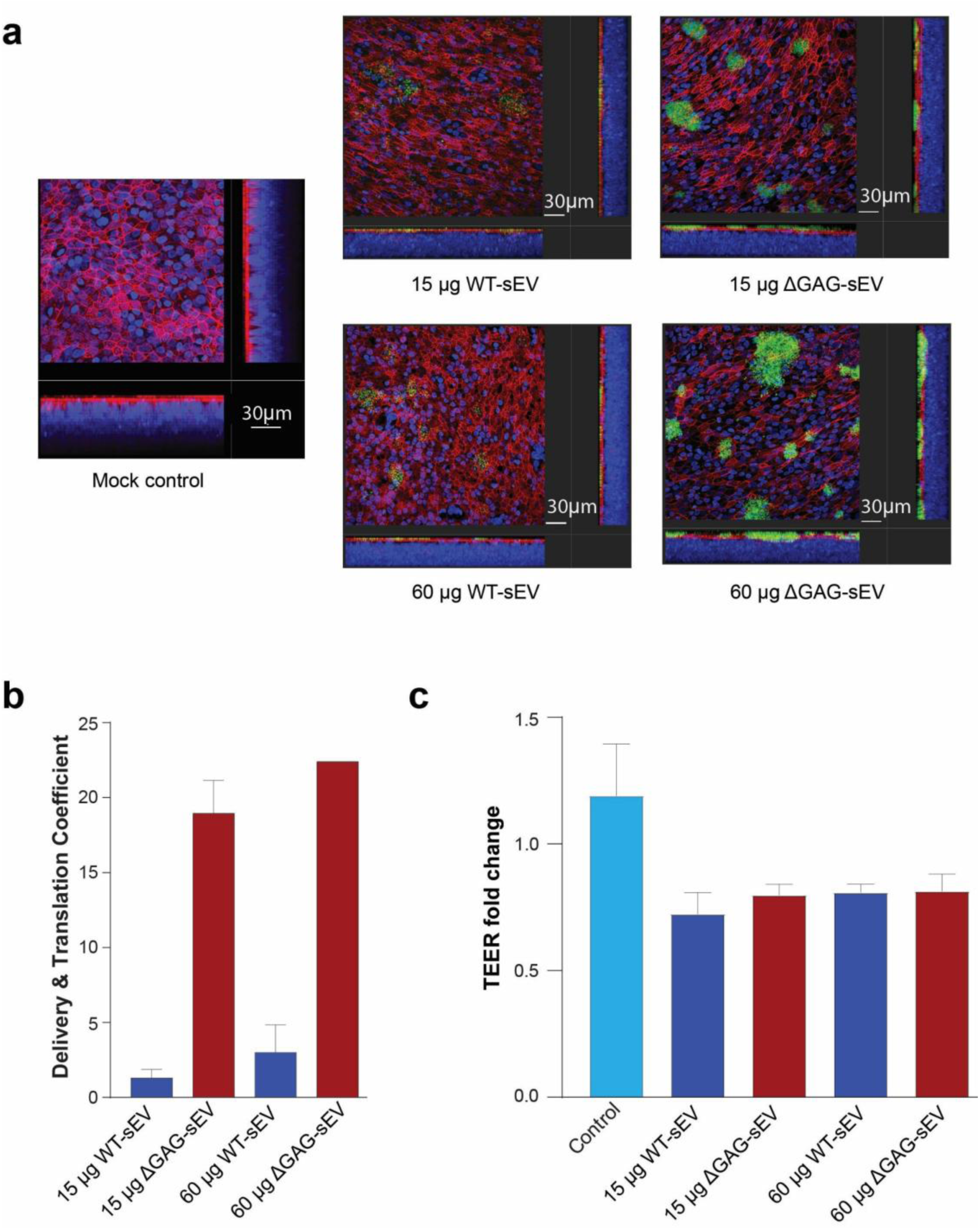
Plasmid DNA delivery by sEVs in an air-liquid interface (ALI) airway epithelial mode. **a,b,** ALI airway models: Differentiated BCi-NS1.1 cells cultured under ALI conditions were treated apically with plasmid-loaded WT- or ΔGAG-sEVs (from a single clone) at two doses (15 µg and 60 µg). GFP expression was assessed 24 h post-treatment by confocal fluorescence microscopy. Representative images (**a**) and quantitative analysis of the DTCs for each group (**b**) are shown for each condition. Images show GFP expressions in green, F-actin in red, and nuclei in blue (DAPI). DTCs signify the total integrated density of GFP per thousand total integrated density of DAPI signal (signifying number of cells in the given ZStack). Data are presented as mean ± s.d. from two replicates, except for the 60 µg ΔGAG-sEV condition (n = 1). **c,** Transepithelial electrical resistance (TEER) was measured to assess epithelial barrier integrity following sEV treatment. Data are presented as mean ± s.d. from four replicates.

## DISCUSSION

Glycosylation is increasingly recognized as a critical determinant of sEV biology, yet the specific contributions of individual glycan classes to vesicle association and functional cargo transfer have remained poorly resolved. Most prior studies probed sEV glycans using enzymatic deglycosylation, but these treatments can alter sEV physicochemical properties and their effects are often partial, variable across vesicle subpopulations, and difficult to attribute to specific glycan classes, complicating mechanistic interpretation [18,22,26,30]. More recent efforts have genetically engineered producer cells to display defined glycan ligands (e.g., sLeX, LeX) on EV surfaces by adding an overexpressed fucosyltransferase. Increased sLeX or LeX structures on EVs were only achieved when an artificial fusion protein with additional glycosylation sites were added in addition to the transferase. The resulting cell-specific targeting focused on exploiting sLeX/LeX-selectin interactions but did not dissect the contributions of endogenous glycan classes to sEV uptake and cargo transfer [31]. Using a genetically defined panel of isogenic HEK293F cell lines in which major glycan biosynthetic pathways are ablated individually, we provide a systematic, quantitative dissection of how sEV- and cell-surface glycans shape vesicle association, internalization, and functional macromolecular transfer.

A central discovery is that GAGs act as dominant inhibitory determinants of sEV association, whether presented on sEVs or on recipient cells, and that removal of GAGs can strongly increase sEV and their cargo delivery including gene transfer and expression by up to 15/30-fold. sEVs lacking GAGs (ΔGAG-EVs) exhibited the highest uptake and the largest increases in functional transfer across cell types and cargo modalities (EV specific dye, proteins, DNA oligonucleotides, siRNA, and plasmid DNA). To minimize potential clonal variability and ensure reproducibility, all uptake assays were performed using sEVs pooled from three independent clones per variant, thereby capturing a generalizable effect. In contrast, sEVs used for ALI experiments were generated from a single randomly selected clone, which may account for the markedly enhanced improvement observed and suggests that selecting optimal sEV-producing clones could further improve delivery efficiency. On the other hand, the strong enhancement in the ALI experiments may be caused by strongly GFP expressing cells for which a >30-fold increase was also observed in the ΔGAG-EV delivery of pMax-GFP plasmid to HEK cells. The breadth of cargo types, and exogeneous as well as endogenous cargo loading, argue that GAG removal enhances a general entry and/or post-entry pathway rather than only facilitating a specific cargo-dependent route. Because functional gene expression from delivered plasmid DNA additionally requires endosomal escape and nuclear entry, our observations are consistent with ΔGAG-sEVs improving one or more downstream steps beyond initial sEV-cell interaction and uptake.

These functional data align with glycan-abundance patterns we previously reported [15]: GAGs are enriched on sEVs relative to parental cell surfaces, whereas GSLs are relatively more abundant on cells. This asymmetric distribution offers mechanistic insights-dense GAG chains on sEVs can sterically hinder protein ligands or create local spatial or electrostatic barriers that reduce receptor engagement, while high cell-surface GSL levels may impose host-side constraints on vesicle binding. Indeed, deletion of the more abundant glycan class on a given membrane (GAGs on sEVs; GSLs on cells) produced larger uptake enhancements than deletion of the less abundant class, highlighting that glycan abundance, not only identity, is a critical determinant of sEV-cell interactions and a practical design parameter for glycoengineering.

Beyond abundance, the chemistry of GAGs likely matters. Heparan sulfate (HS) and chondroitin/dermatan sulfates (CS/DS) vary in sulfation motifs and backbone structure and engage different protein partners. Our genetic ΔGAG approach stops the build-up of the 4 sugar units linker glycan to which the long GAG chains are linked at the first sugar thus preventing all HS, CS and DS GAGs from forming simultaneously. Understanding which GAG subtypes (e.g., HS vs CS/DS) and which sulfation patterns drive the ΔGAG phenotype will require more targeted glycoengineering and detailed glycoanalytics [20,32–34]. Notably, removal of sialic acids (ΔSA-sEVs) produced a smaller but consistent increase in cellular uptake, in line with previous reports using enzymatic approaches [22,35]. Although ΔSA-sEVs displayed the least negative zeta potential, ΔGAG-sEVs showed markedly superior uptake and cargo delivery performance, indicating that surface charge alone cannot account for the observed improvements (Fig. S1). The fact that ΔGAG-sEVs and ΔNG-sEVs showed a comparable zeta potential, the most negative of all glycoengineered sEV and closest to WT-sEV, further supports a more complex phenomena than charge.

The pairing of glycoengineered sEVs with glycoengineered recipient cells in a matrix design revealed unexpected interaction effects underscoring the power of our isogenic cell-based glycoengineering platform. ΔGAG-sEVs achieved maximal uptake in ΔGSL and ΔSA recipient backgrounds rather than in ΔGAG cells, implying compensatory and cooperative contributions across glycan classes. These multivalent effects underscore the complexity of carbohydrate-mediated recognition and caution against interpreting single-glycan perturbations in isolation. They also point to the potential for rational combinatorial glycoengineering, tuning both vesicle and target membranes, to maximize uptake and functional transfer for specific therapeutic contexts.

Lipid nanoparticles (LNPs) and viral vectors are currently the most used platforms for clinical gene delivery, yet both exhibit significant limitations [36,37]. Viral vectors are associated with immunogenicity preventing repeat administration, risks of insertional mutagenesis, and restricted capacity for type and cargo size [36,38,39]. Conversely, LNPs tend to accumulate in the liver and face challenges related to toxicity [40,41]. GE-sEVs combine native membrane complexity with programmable surface chemistry, potentially enabling diverse cargo types and improved biocompatibility. The large increase of up to 15/30-fold for plasmid delivery and gene expression makes ΔGAG-sEVs a promising approach for drug delivery and gene therapy.

In summary, our work establishes glycan abundance and composition as tunable parameters that can be engineered to substantially improve sEV-cell association and macromolecular transfer. Removal of GAGs emerges as a robust strategy to enhance functional transfer across cargo types and cell models and for improved sEV drug delivery and gene therapy approaches. Systematic glycoengineering therefore opens a new frontier for creating optimized, programmable vesicle therapeutics across gene delivery, immunotherapy, and regenerative medicine.

## ACKNOWLEDGMENTS

This work was supported by Novo Nordisk Foundation grants NNF19SA0056783, NNF19SA0057794, NNF20SA0066621, and NNF24OC0090768 to S. G., and by Novo Nordisk Foundation grant NNF19OC0056411 and a grant from the John and Birthe Meyer Foundation to H. K. J.

## AUTHOR CONTRIBUTIONS

S.G. supervised the project and collected financial support. W.T. and S.G. conceptualized, planned experiments, analyzed the data and wrote the manuscript. W.T. engineered and generated cells, sEV, and performed most of experiments. J.C., and J.P.A. helped cell culture, sEV isolation and characterization; A.L.B. and L.E.P. helped data analysis; A.J.F.M., A.M.R, H. K. J and S.M performed and financially supported ALI experiments.

---

**Fig. S1:**
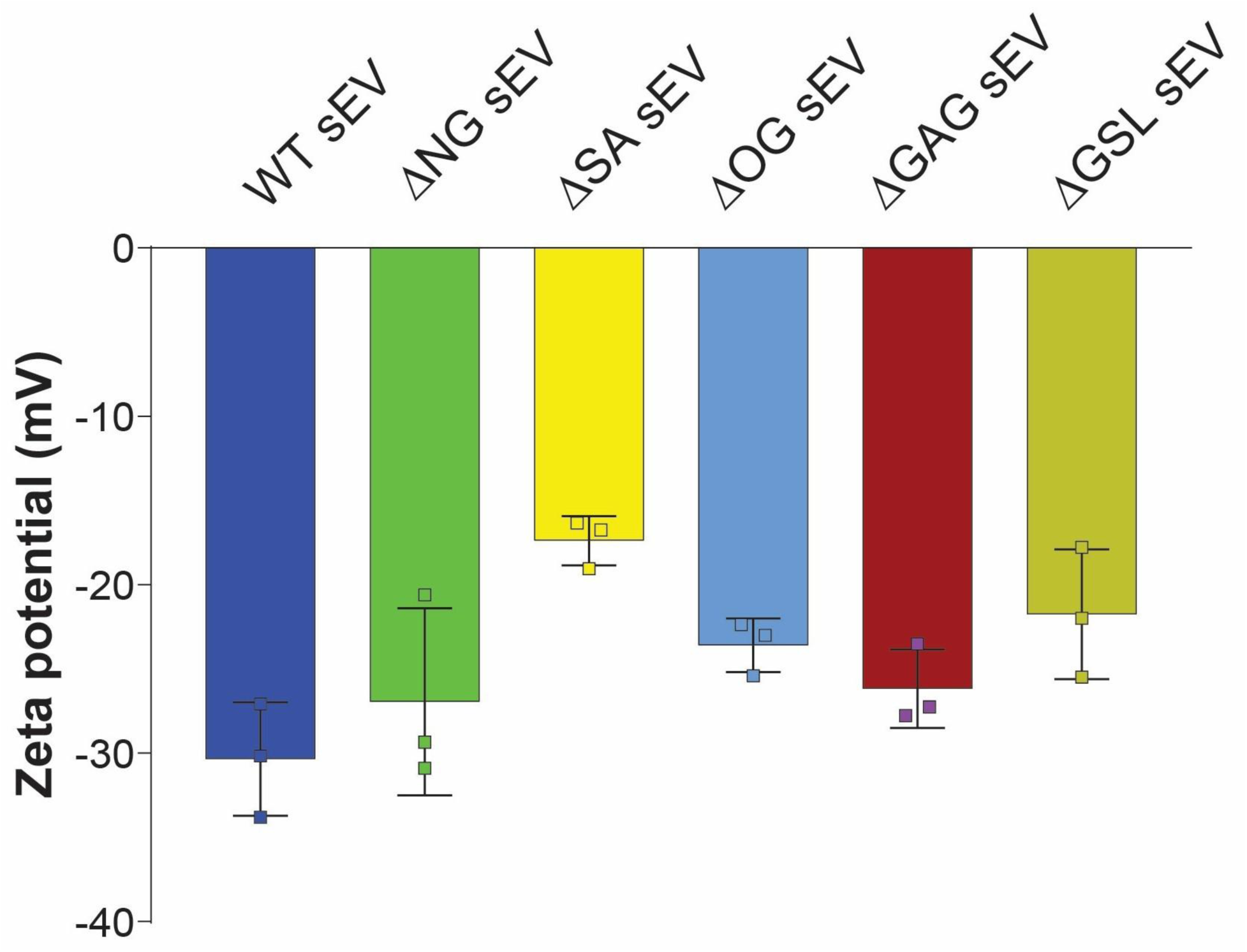
Zeta potential characterization of glycoengineered sEVs. Zeta potential was measured using the Particle Metrix ZetaView NTA system. Three independent clones of each variant and three separate batches of HEK293F WT cells were measured. Data are presented as mean ± s.d. from three replicates.

